# Delivery mode impacts newborn gut colonization efficiency

**DOI:** 10.1101/2020.01.29.919993

**Authors:** Caroline Mitchell, Larson Hogstrom, Allison Bryant, Agnes Bergerat, Avital Cher, Shawna Pochan, Penelope Herman, Maureen Carrigan, Karen Sharp, Curtis Huttenhower, Eric S. Lander, Hera Vlamakis, Ramnik J. Xavier, Moran Yassour

## Abstract

Delivery mode is the variable with the greatest influence on the infant gut microbiome composition in the first few months of life. Children born by Cesarean section (C-section) lack species from the *Bacteroides* genus in their gut microbial community, and this difference can be detectable until 6-18 months of age. One hypothesis is that these differences stem from lack of exposure to the maternal vaginal microbiome, as children born by C-section do not pass through the birth canal; however, *Bacteroides* species are not common members of the vaginal microbiome, thus this explanation seems inadequate. Here, we set out to re-evaluate this hypothesis by collecting rectal and vaginal samples before delivery from 73 mothers with paired stool from their infants in the first two weeks of life. We compared microbial profiles of infants born by planned, pre-labor C-section to those born by emergent, post-labor surgery (where the child was in the birth canal, but eventually delivered through an abdominal incision), and found no significant differences in the microbiome between these two groups. Both groups showed the characteristic signature lack of *Bacteroides* species, despite their difference in exposure to the birth canal. Surprisingly, this signature was only evident in samples from week two of life, but not in the first week. Children born by C-section often had high abundance of *Bacteroides* in their first few days of life, but these were not stable colonizers of the infant gut, as they were not detectable by week two. Finally, we used metagenomic sequencing to compare microbial strains in maternal vaginal and rectal samples and samples from their infants; we found evidence for mother-to-child transmission of rectal rather than vaginal strains. These results challenge birth canal exposure as the dominant factor in infant gut microbiome establishment and implicate colonization efficiency rather than exposure as a dictating factor of the newborn gut microbiome composition.

## Main text

Infants begin life with a relatively simple gut microbial community and acquire a more complex, adult-like microbiome over the first three years of life^1–4^. While multiple factors influence composition and dynamics during the first year of life, delivery mode is one of the most significant^5,6^. Infants born by Cesarean section (C-section) have significantly different early gut microbial features compared to vaginally delivered infants, including lower diversity and richness^2,7^ and lower prevalence of colonization with *Bacteroides* species^1,5–9^. This C-section microbial signature is detectable to at least 6 months of age despite subsequent effects of breastfeeding on the infant microbiome^1,4,5^.

Pioneering studies demonstrated the difference between delivery modes on early microbiome development, but did not identify the underlying causes of this difference. It is widely believed that the infant gut is seeded with microbes from the mother during labor, particularly from the maternal vaginal microbiome. The data supporting these hypotheses, however, are limited. Some studies have relied on a single infant stool sample collected around birth^1,2,8,10–12^, examined a small number of 20-30 families^7,8,13–15^, confounded vaginal microbiome exposure by combining pre- and post-labor C-section groups (infants born by post-labor C-section are exposed to the maternal vaginal microbiome, but eventually extracted by C-section)^2,4,6,11^, or excluded post-labor C-sections altogether^8,12,14 2,4,6,11^.

Infants born by post-labor C-section provide a unique opportunity to study the impact of traveling through the birth canal on bacterial transmission from mother to infant. Comparing the microbiomes of infants born by pre- and post-labor C-sections will more definitively address the importance of exposure to vaginal microbes independent of mode of exit. Furthermore, studying mother-to-infant microbial transmission requires a comparison of the microbes found in both mothers and infants at the strain-level, which can only be done using deep metagenomic sequencing, as we recently reported^15,16^.

Here, we examine the sources and timing of delivery mode-associated microbial signatures. We established a birth cohort to collect multiple newborn and maternal microbiome samples from the first two weeks of life. Using metagenomic sequencing, we compared the gut microbiome of newborns across delivery modes and found that (1) the vaginal delivery microbial signature is not simply the result of exposure to birth canal microbes, as children born by post-labor C-section had a microbial signature resembling those born by pre-labor C-section; (2) the lack of *Bacteroides* species in the guts of infants delivered by both pre- and post-labor C-sections only appears in the second week of life, as many children had detectable *Bacteroides* in their first week of life that later disappeared; and (3) mother-to-child bacterial transmission events occur mostly in vaginally delivered children, and the maternal source is rectal rather than vaginal.

To study microbiome acquisition immediately following birth, we collected maternal and newborn samples from 73 families admitted to the Massachusetts General Hospital Labor and Delivery unit. We enrolled 38 children born vaginally, 19 children born by scheduled, pre-labor C-section, and 16 children born by non-elective C-section following labor. We collected maternal rectal and vaginal samples less than 24 hours prior to delivery to evaluate potential maternal microbial sources at the time of delivery. In addition, we collected daily stool samples from the newborns while they were at the hospital (2-4 days) and one stool sample at two weeks of age, after hospital discharge (see **Fig. 1A** and **Supplemental Table 1,2** for characteristics of the cohort). Neonatal stool samples often have low bacterial biomass and varying sample volumes, which motivated our collection of multiple samples from the first week of life to obtain a more complete picture of initial microbial exposure and colonization. We performed 16S rRNA gene sequencing on infant samples and metagenomic sequencing on both maternal and infant samples.

**Figure 1:**
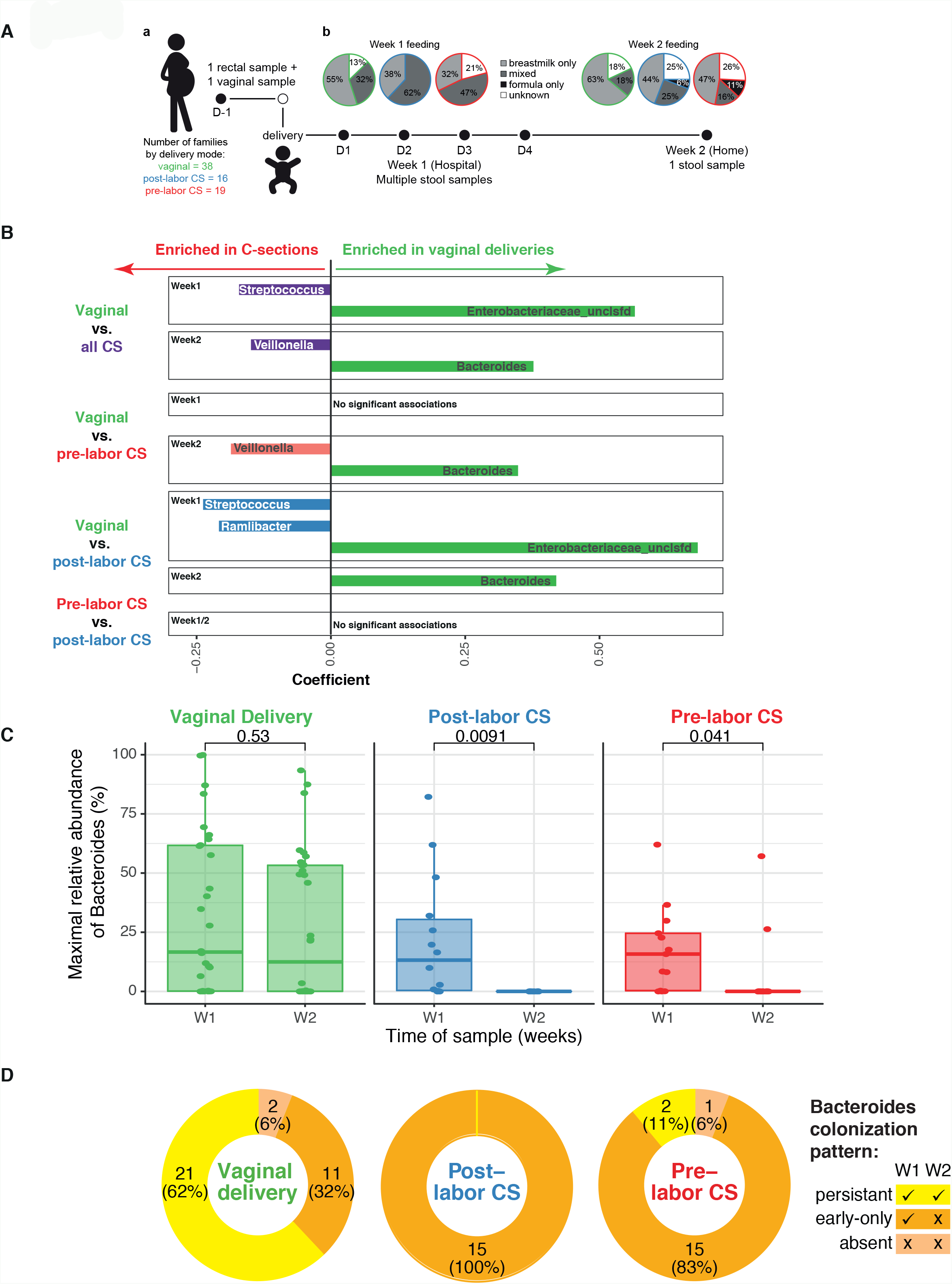
(A) Cohort description. Women presenting to Labor and Delivery at term (≥ 37 weeks) with singleton gestation were enrolled, and had a rectal and vaginal sample collected prior to delivery. Infants had stool collected daily while in hospital, and sent a single stool sample from home at 2 weeks of life. Feeding practices were abstracted from inpatient charts for week 1 and obtained from parent questionnaires for week 2. **(B) Impact of delivery mode on early life microbial composition.** MaAsLiN was used to identify taxa that were over-represented in vaginally delivered vs. C-section delivered infants, vaginally delivered vs. pre- or post-labor C-section, and pre- vs. post-labor C-section (using the results of 16S rRNA sequencing). Analyses were adjusted for mode of feeding in the week of interest. **(C)** Maximal abundance of *Bacteroides* in week 1 or week 2 samples was compared between vaginal delivery, pre- or post-labor C-section delivery using t-test. **(D)** The *Bacteroides* colonization phenotype was assigned based on detection at > 1% in week 1 samples only (*early-only*), both week 1 and week 2 (*persistent*), or neither (*absent*).

To determine how delivery mode affects acquisition and persistence of bacterial taxa, we stratified samples into week 1 (multiple time points per infant) and week 2 (one time point per infant). Due to gut microbiome variability in the first few days of life^16^, we calculated the maximum abundance for each taxon across the multiple week 1 samples from each infant samples, in order to consider all species that the infant was exposed to in the first week of life. We compared genus-level gut microbiome profiles across delivery modes using multivariate linear regression with infant feeding as a confounding variable.

Surprisingly, the well-established absence of *Bacteroides* species in the gut microbiome of most infants born by C-section was not evident in the first week of life (**Fig. 1B, C)**. We found that 19/35 (54%) infants born by pre- or post-labor C-section had detectable levels of gut *Bacteroides* species (≥0.5%) during their first week of life. However, nearly all (94%) infants showed no detectable levels in their second week of life (confirmed by both metagenomics and 16S rRNA gene sequencing). Consistent with the literature, we found that at week 2, infants born via C-section (both pre- and post-labor) were much less likely to be colonized by *Bacteroides* relative to infants born vaginally (**Fig. 1B, C**). Importantly, we used maximal abundance with 16S rRNA gene sequencing for its sensitivity to detect low abundance taxa; the results are also mirrored in the metagenomic data (**Supplemental Figure 1A**). Because breastfeeding rates and exposure to formula differed by delivery mode, we adjusted for infant feeding in all analyses.

Notably, we found no significant differences between the microbiomes of infants born by pre- and post-labor C-sections, suggesting that vaginal exposure may not be a driving force in *Bacteroides* colonization (**Fig. 1B**). We confirmed these two results concerning the *Bacteroides* signature (the difference between weeks 1 and 2 and lack of difference between pre- and post-labor C-section) by re-examining published datasets. By analyzing the raw data from a published cohort^10^, we observed the same pattern of early *Bacteroides* colonization and later disappearance in some children delivered by C-section **(Supplemental Figure 1B**). This difference between week 1 and 2 had not been identified or highlighted by the authors. The early presence of *Bacteroides* species suggests that their later absence cannot be attributed to lack of exposure^13,17^.

To facilitate analysis below, we grouped infants according to the pattern of *Bacteroides* colonization in weeks 1 and 2. We defined *Bacteroides* colonization as *persistent* if it was detected in week 1 and 2 samples, *early-only* if it was detected in week 1 but not week 2, and *absent* if it was not detected in neither week 1 or 2. A small number of children (3) had the *absent* colonization pattern, and no infants in our study were only colonized at week 2. The vast majority of children born by C-section (91%) have an *early-only* colonization pattern of *Bacteroides*, whereas most vaginally delivered children (62%) have the *persistent* colonization pattern (**Fig. 1D**).

We wondered whether maternal factors other than delivery mode might explain the differential *Bacteroides* colonization patterns. However, we saw no difference across delivery modes in maternal demographic characteristics, type of anesthesia received, and prevalence of medical co-morbidities, which suggests that these characteristics were not significant confounders **(Supplemental Table 2**). In addition, all women undergoing C-section receive antibiotics prior to incision, and we wanted to confirm that the differential colonization pattern across delivery modes was not due to the antibiotic treatment. Some of the women who delivered vaginally were also given antibiotics—for example, if they tested positive for group B Streptococcus (GBS). To test the hypothesis that antibiotic treatment was associated with the loss of *Bacteroides* in week 2, we examined the relative abundance of *Bacteroides* in vaginally-delivered children at week 2, but found no effect of maternal antibiotics at labor on the *Bacteroides* colonization pattern (**Supplemental Figure 1C**).

We next explored what might cause the loss of *Bacteroides* in week 2. We first compared infant characteristics across the *Bacteroides* colonization patterns. Detection of individual *Bacteroides* species was similar between vaginally and C-section delivered infants in week 1 samples, suggesting no species differences in initial colonization. While delivery mode was significantly associated with *Bacteroides* colonization, infant characteristics such as exposure to formula in the hospital, breastfeeding at week 2, and infant receipt of antibiotics were not (**Supplemental Table 3**). While not significantly associated with *Bacteroides* colonization, exposure to formula in the first week of life was higher after C-section delivery (51% vs. 25%, p = 0.02), so analyses were controlled for this variable. We also compared the duration of vaginal exposure in infants exposed to labor (vaginal or post-labor C-section deliveries): neither the length of second stage of labor (time between full dilation and birth) nor time between membrane rupture and delivery was associated with persistent vs. early *Bacteroides* colonization in infants born vaginally or by post-labor C-section (not shown), indicating, again, that the persistence of *Bacteroides* colonization is not a result of vaginal exposure.

We next wondered whether maternal factors influenced the infant *Bacteroides* colonization patterns. We examined the relationship between maternal microbial communities and the infant *Bacteroides* colonization phenotype. We found no difference across delivery modes in the relative abundance of maternal rectal *Bacteroides* between the colonization phenotypes (not shown). Only a single maternal vaginal sample had detectable (>0.5%) *Bacteroides* colonization, which could not account for the *Bacteroides* transmission (“V3”; see **Supplementary Figure 2A**). Some authors have suggested that the differences seen in the infant gut microbiome due to delivery mode point to underlying differences in women who have C-sections compared to those who deliver vaginally. Maternal rectal *Bacteroides* abundance was not different between the three delivery modes, suggesting that differences in infant colonization were not due to an association between maternal colonization and delivery mode (not shown).

**Figure 2:**
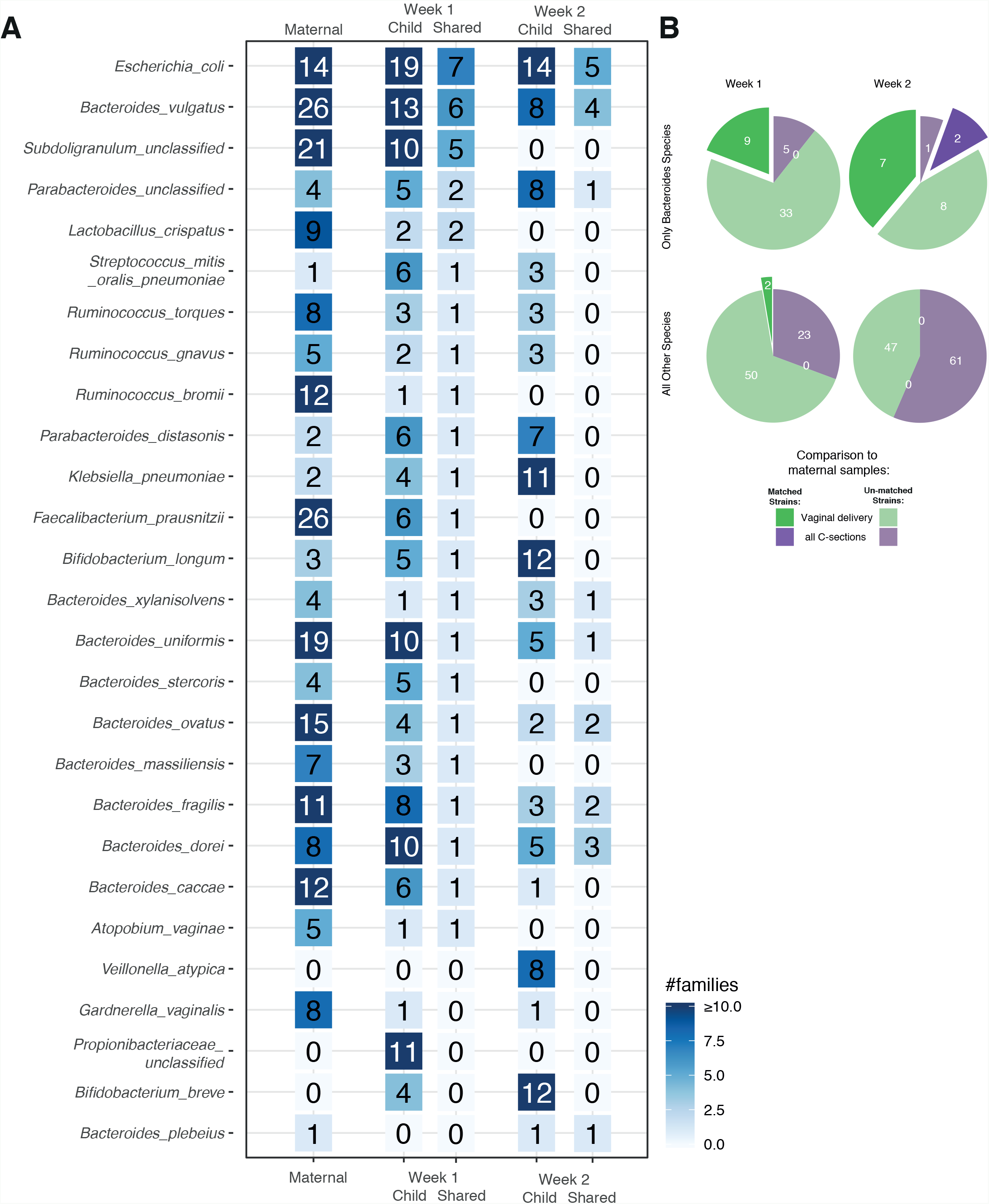
Shared species and strain between maternal rectal and infant stool samples. Using metagenomic sequencing of infant stool and maternal rectal samples,**(A)** the number of families with a given species detected (>= 1%) in either mother, child or both in week 1 or week 2 was calculated. For families with shared species, **(B)** strain level comparison was made using species-specific marker genes to identify whether mother and infant shared the same strain.

Finally, we hypothesized that the later loss of Bacteroides may be due to competition within the infant gut. To address this question, we compared week 1 samples from all of the C-section infants to those born vaginally to identify additional taxa that might be associated with the delivery mode and the loss of *Bacteroides* in week 2. Interestingly, we found an increase in the genus *Streptococcus* in C-section born children in the first week of life, prior to the disappearance of *Bacteroides* (**Fig. 2B, Supplemental Figure 1D**). When we compared vaginally delivered infants by *Bacteroides* colonization phenotype, we found that in week 2 samples the abundance of *Streptococcus* species was negatively associated with persistent *Bacteroides* colonization (**Supplemental Figure 1E**), suggesting that competition between these species may be a driving force in the loss of *Bacteroides* in the second week of life.

We next asked, for children in whom we observe *Bacteroides* in week 1 samples, what is the source of these strains? Specifically, are the infant strains shared with their mothers? It is possible that the source of the *Bacteroides* strains plays a role in its persistence in an infant’s gut. The data above suggest that *Bacteroides* colonization is not dependent on whether an infant was exposed to the birth canal, but rather whether the infant was extracted by C-section or delivered vaginally. Therefore, for analysis of the source of infant colonization, we considered all C-sections together as a single group. Because exiting the birth canal brings the infant in proximity to the maternal rectum, we hypothesized that during a vaginal delivery, the infant is exposed to bacterial strains from the maternal gut microbiota, and maternal *Bacteroides* strains may have features that favor persistence in the infant gut.

To obtain species-level resolution and be able to ask strain-level questions, we used taxonomic profiling from metagenomic data. As the infant and adult gut harbor different microbial communities, we are limited in this analysis to species that are shared within enough families (**Fig. 2A**). Among 9 vaginally delivered infants with detectable *Bacteroides* in week 1 for which we also had sufficient coverage of their mother’s rectal samples, 8 (88%) shared at least one *Bacteroides* species with their mother compared to 3/6 (50%) C-section delivered infants, suggesting indeed that delivery mode (and perhaps exiting the birth canal specifically) increases infant colonization by maternal species.

We next turned to identify matched strains between mother and infant species that might suggest a possible transmission event, using single nucleotide variants extracted from the metagenomic reads (see **Methods**). Among the 9 of 42 *Bacteroides* strains in week 1 that matched a strain found in the mother, all occurred in vaginally-delivered children (**Fig. 2B**). None of the 5 *Bacteroides* strains found in children born by C-section were shared with their mothers. In week 2, 9/18 (50%) infant *Bacteroides* strains matched their maternal strains, and of those, the vast majority 7/9 (78%) were identified in infants born vaginally. Overall, across all species and all time points, 18/20 (90%) of matched maternal-infant strains were found in infants born vaginally (**Figure 3B**).

Finally, it has been suggested that vaginally delivered infants acquire *Lactobacillus* species from their mothers as they pass through the birth canal^13^. Despite the high abundance of *Lactobacillus* in our vaginal samples (**Supplemental Figure 2A**), we did not observe higher abundance of *Lactobacillus* species in the gut microbiome of children born vaginally. Only two infants had *Lactobacillus* species detected at week 1 samples (above 2%), both vaginally delivered. In both cases, maternal samples had a high proportion (>90%) of the same *Lactobacillus* species. If we extend this analysis to additional common vaginal microbes, among 29 species found at >2% in any maternal vaginal samples, only 5 were shared between mothers and infants and each appeared in only a single family (**Supplemental Figure 2B**), suggesting that transmission of maternal vaginal species to the infant is infrequent and/or in low abundance.

To summarize, we sequenced microbiomes from 73 newborns and their mothers close to the time of delivery to determine the impact of delivery mode on the origin of the infant gut microbiome.

Our comparison between children born by post-labor C-section and vaginal delivery highlights a canonical C-section microbial signature in all types of C-sections, suggesting the major factor driving the association between birth mode and infant microbiota is not entering the birth canal, but rather exiting through it. The similarity of infant gut communities between infants delivered by C-section before or after labor is consistent with findings in the Baby Biome Study from the United Kingdom^18^ which sampled infants once in the first week of life. We did not find evidence of vaginal microbiota transmission from mothers to their vaginally delivered infants, and very few infants had detectable levels of *Lactobacillus* (the common vaginal microbiome member) in their gut community, also consistent with Baby Biome Study findings^18^. Indeed, the strongest strain-level evidence for bacterial transmission at birth is found in vaginally delivered children but stems from the maternal rectal source. Our results suggest that the establishment of the initial infant gut microbial community requires more than topical exposure to maternal vaginal microbes.

Multiple infant samples collected during the first week of life enabled us to question the timing of the canonical C-section microbial signature, namely the depletion of *Bacteroides* species in the first 6-18 months. To our surprise, we found that 51% of infants born by pre- or post-labor C-section had detectable levels of gut *Bacteroides* species during their first week of life, which disappeared in their second week of life (confirmed by both metagenomics and 16S sequencing). These results raise interesting questions: First, why do these *Bacteroides* disappear? Second, from where are they acquired? In 18/65 (28%) infant samples where *Bacteroides* species were detected, we were able to identify matched strains in the maternal rectal swabs that appear to be the source of *Bacteroides* species; but more often than not, we were unable to identify the source strain from mother vaginal or rectal samples.

Our data suggest that the origin of infant microbial differences may lie in both the source of the microbes and the infant environment that they are colonizing. A well-characterized cohort of vaginally delivered infants suggested that infant “seeding” likely originated from several maternal sites, and that the ability of those microbes to persist might be related to the relative fitness of a given isolate for the gut environment^15^. Our findings support the hypothesis that a surgical delivery alters the receptivity of the infant gut environment. The disappearance of *Bacteroides* species in the second week of life from infants born via C-section suggests either the lack of a supporting factor or the introduction of an antagonist during the surgical process. We reviewed charts to identify any exposure to formula during the delivery hospitalization, and we found a much higher rate of formula exposure in C-section delivered infants, although most infants in our cohort were primarily breastfed. Formula feeding may be one of several factors that contribute to a difference in gut microbiota of infants born via C-section. In the CHILD cohort, even a single exposure to formula during the delivery hospitalization was associated with lower prevalence of *Bifidobacteriacea* and higher prevalence of *Enterobacteriaceae* at 3 months ^19^. We adjusted for feeding practices in our analyses, but larger cohorts are needed to further study these influences.

Although we did not identify a specific microbe in early life samples that was negatively associated with establishing persistent *Bacteroides* colonization, we did identify higher prevalence of *Streptococcus* species in the second week of life among those infants who lost *Bacteroides* colonization. The striking negative correlation between *Bacteroides* and *Streptococcus* could be due to *Streptococcus* species filling a niche that was vacated, or to direct competition between the two genera. Members of the *Streptococcus mitis* group can produce hydrogen peroxide, which inhibits growth of other oral microbes^20,21^. *Streptococcus oralis* can inhibit pathogen growth and biofilm formation^22^. Either of these mechanisms could account for the negative correlation between *Streptococcus* and *Bacteroides*.

Subsequent studies examining additional microbial sources and the infant gut metabolic environment will be critical for understanding the origin and persistence of these species. Although feeding practices play a significant role in determining the composition of the infant gut microbiota^2,19^, understanding the earliest microbial communities and their determinants is an important component of designing microbiome-related interventions to improve infant health. Part of the challenge in early life samples is the low biomass of the stool samples. Fortunately, improved methods for profiling low-biomass samples^23,24^ make it increasingly feasible to study the strain-level infant gut colonization at the very early stages of life.

Our results demonstrate that maternal seeding of the infant gut microbiome can occur, but is not a straightforward transfer of the dominant maternal strains of a given species. We also show that the impact of C-section delivery on the infant gut community is a delayed effect, and thus as much an influence on the infant gut environment as on the source of originating bacteria. These findings demonstrate the complexity of establishing a microbial community and our lack of understanding of how obstetric care influences neonates.

## Materials and Methods

### Study participants

Women with singleton gestation, presenting for delivery at >= 37 weeks of gestation to the Massachusetts General Hospital Labor & Delivery unit in Boston, MA were enrolled in this prospective cohort study. Participants were excluded if they had known HIV infection, fetal congenital anomaly, gestational age < 37 weeks, maternal age < 18, planned to give up the infant for adoption, or were a gestational carrier. If an enrolled family had subsequent admission of the infant to the NICU or special care nursery at any time after delivery, that family was excluded. At enrollment, maternal participants had vaginal and rectal swabs collected. Infant stool was collected by parents from the diaper once daily during the delivery hospitalization, into tubes containing ethanol preservative. Study staff collected infant stool samples from parents each day. At 2 weeks of life, a sample collection kit was sent to parents, who returned another infant stool sample (also in ethanol) within 24 hours of collection, via a delivery service. Participants answered a short questionnaire about infant health status and mode of feeding, which was returned with the 2 week stool sample. All stool was placed in the freezer within 2 days of collection. Participant and infant medical records were abstracted to obtain information about mode of delivery, receipt of antibiotics, reported infant feeding, duration of labor stages, obstetric complications, and health history.

### DNA Extraction and sequencing

Vaginal and rectal swabs were eluted in 400uL of saline, vortexed for 1 minute and then spun at 10,000 xg for 10 minutes. The cell pellet underwent DNA extraction using the MoBio Power Soil kit (Qiagen Inc, Waltham MA). Stool samples underwent DNA extraction using chemical and mechanical lysis with magnetic bead-based purification via the Chemagic MSM I with the Chemagic DNA Blood Kit-96 from Perkin Elmer. Prior to extraction on the MSM-I, TE buffer, lysozyme, proteinase K, and RLT buffer with beta-mercaptoethanol were added to each stool sample. DNA samples were quantified using a fluorescence-based PicoGreen assay.

### Sequence processing and analysis

A total of 426 samples from 84 families underwent sequencing. Whole-genome shotgun (WGS) sequencing was performed on the Illumina NextSeq platform with 100 bp paired-end reads. Reads pairs with fewer than 60 observed bases on either read were excluded from downstream analysis. Samples with fewer than 500,000 raw reads were excluded. Read pairs that were attributed to human genomic DNA were removed using KneadData Tool, v0.5.1 using the hg19 human reference genome. Samples with fewer than 100,000 non-human reads were also excluded from downstream analysis. To taxonomically profile the bacterial DNA sequences observed in the samples, MetaPhlAn 2.0 was used to align reads to the MetaPhlAn 2.0 unique marker database. Samples that were fully unclassified by metaphlan were excluded. Samples were also excluded if there were 100 or fewer reads mapping to the species-specific marker genes of a given taxa that was reported at >50% abundance. For inclusion in downstream analysis, samples coming from a given family had to pass the quality measures above and meet at least one of the following family criteria: a) two or more child samples were available in week 1, b) One or more child samples for week 1 were available and one child sample at week 2, or c) one or more child samples and one or more mother sample available. A subset of 73 families met one or more of these criteria for inclusion. There were 71 infant stool samples from this subset that had sufficient DNA and underwent 16S rRNA sequencing of the V3-V4 region.16S rRNA gene sequencing was performed essentially as previously described^4^. Taxonomy was assigned using version 1.8.0 of Qiime^25^ and the Greengenes reference database of OTUs^26^. The number of reads for both 16S and WGS methods was not different across delivery modes.

### Mother-child strain transmission analysis

We tabulated the species that were observed most frequently in both mother and child samples of the same family (**Fig. 2A**). To examine families for possible shared strains within these species, we applied an approach we recently developed to examine sub-species variation between samples^16^. Briefly, coverage overlap events were identified in mother-child pairs where metagenomic reads mapped to the same segment of species-specific marker genes of various species. In these overlap regions, sites of single nucleotide variation (SNV) were tabulated across the set of marker genes for a given species. Samples with fewer than 1000 reads mapping to species-specific marker genes were not considered for this analysis.

We first identified “major-major” nucleotide match events at positions where the reads coming from the mother sample contained a most frequent base that matched those seen most frequently in child reads. The frequency of these events were used to estimate a shared dominant strain of a given species found both mother and child samples. We also examined ‘‘MotherMinor-ChildMajor’’ match events at positions where the second most frequent nucleotide found in the mother’s reads matched the most frequent nucleotide found in the child’s reads. The frequency of these events were used to estimate shared secondary strains of a given species found both mother and child samples.

In addition to testing for sites of shared variation between mother-child samples from the same family, we also quantified shared variation occurring within samples of unrelated subjects. The relative frequency of match and mismatch events were attributed to either dominant or secondary strains. In both cases, match rates coming from mother-child pairs of the same family were converted to Z-scores using a null distribution created by comparing sample pairs from unrelated families^16^.

#### Statistical analysis

A *Bacteroides* colonization phenotype of early-only (week 1 only), persistent (both week 1 and 2) or absent was assigned using 16S rRNA sequencing results, setting “present” at a threshold of 0.5%. Associations between maternal and delivery characteristics and this phenotype were examined using chi-square or ANOVA. Associations between individual taxa and delivery mode, adjusting for infant feeding, were evaluating using MaAsLin^27^.

### Data Availability

All metagenomic data are available in NCBI Sequence Read Archive as BioProject PRJNA591079. Full metadata is available in Supplementary Table 1.

### Approval for human patient research

Human patient research in the OriGiN cohort was reviewed and approved by the Partners Human Research Committee (ref. 2015P000460/PHS). Each mother signed two informed consent forms (one for her and one for her child) prior to participation.

### Figures

All figures were generated with the R ggplot2 package (version 3.1.0). Box plots were generated with geom_boxplot default parameters, such that the box represents 25-75 percentile, and the whiskers from the box to the largest/smallest value no further than 1.5 * IQR from the box (where IQR is the inter-quartile range, or distance between the first and third quartiles).

## Acknowledgements

We thank T. Poon, S. Steelman and C. Nusbaum (Broad Institute) for help in sequence production and sample management; V. Subramanian and T. Arthur for valuable experimental support and helpful discussions; The nurses, midwives and clinicians at the labor and delivery department at Massachusetts General Hospital for their enthusiastic collaboration during enrollment; The families that agreed to enroll and donate samples. Funding: CM was supported by the Vincent Memorial Research Funds; E.S.L. was supported by National Human Genome Research Institute grant 2U54HG003067-10; R.J.X. was supported by funding from JDRF; CCFA, NIH grants R01 DK092405, and P30 DK043351; M.Y. was supported by NIH grant DK113224-01, and a BroadIgnite grant.

## Supplemental Figures

**Supplemental Figure 1: (A)** Maximal relative abundance of *Bacteroides* in week 1 or week 2 infant stool samples, as measured by metagenomic sequencing, compared between vaginally delivered infants and those born after pre- or post-labor C-section. **(B)** Maximal relative abundance of *Bacteroides* by 16S sequencing in delivery or week 6 infant stool samples from 81 infants enrolled by Chu et al. in Houston, TX. **(C)** Comparison of maximal abundance of Bacteroides in week 1 vs. week 2 infant stool samples within vaginally delivered infants whose mothers did or did not receive antibiotics during labor, or infants delivered by pre- or post-labor C-section (all of whose mothers received antibiotics prior to delivery). **(D)** Maximal relative abundance of *Streptococcus* measured by 16S rRNA sequencing compared between vaginally delivered and C-section delivered infants at week 1 or week 2. **(E)** Relative abundance of infant *Streptococcus* in week 2 across Bacteroides colonization patterns.

**Supplemental Figure 2: (A)** Genus-level microbiome composition of maternal vaginal swabs collected prior to delivery, using metagenomic sequencing. **(B)** Shared species between maternal vaginal samples and infant stool samples (as in Figure 2A, only for the vaginal samples).

**Supplemental Table 1:** Full clinical metadata for the 73 families included in the analysis (Excel file).

**Supplemental Table 2:** Metadata statistics for the 73 families included in the analysis

|  | Vaginal | Post-labor | Pre-labor | P value* |
| --- | --- | --- | --- | --- |
| Age | 33 ± 5 | 31 ± 5 | 38 ± 4 | < 0.001 |
| White race | 29 (76%) | 11 (69%) | 18 (95%) | 0.19 |
| BMI | 28 ± 8 | 28 ± 5 | 28 ± 6 | 0.24 |
| Gestational diabetes | 0 (0%) | 1 (6%) | 1 (6%) | 0.32 |
| Hypertension | 6 (16%) | 2 (13%) | 1 (5%) | 0.52 |
| Asthma | 7 (18%) | 3 (19%) | 2 (11%) | 0.72 |
| Chorioamnionitis | 7 (18%) | 3 (19%) | 0 (0%) | 0.13 |
| Maternal antibiotics | 18 (47%) | 16 (100%) | 19 (100%) | < 0.001 |
| Meconium | 12 (32%) | 6 (38%) | 1 (5%) | 0.05 |
| Birthweight | 3515 ± 416 | 3331 ± 421 | 3687 ± 528 | 0.07 |
| Infant antibiotics | 6 (16%) | 4 (25%) | 0 (0%) | 0.09 |
| Any formula in hospital | 9 (24%) | 9 (56%) | 9 (47%) | 0.04 |
| Breastfeeding (hospital discharge) | 27 (75%) | 7 (44%) | 10 (53%) | 0.06 |
| Exclusive breastfeeding (2 weeks) | 24 (63%) | 7 (44%) | 9 (47%) | 0.63 |
| * chi square or ANOVA |  |  |  |  |

**Supplemental Table 3:** *Bacteroides* colonization was characterized as *absent* (not detected at week 1 or 2), *early-only* (detected at week 1 only), *late* (detected at week 2 only) or *persistent* (detected at both week 1 and week 2). A total of 67 families had data from 16S sequencing for both week 1 and week 2 which allowed this classification. Detection was defined as a relative abundance of 0.5%. Groups were compared by chi-square or ANOVA.

|  | *Absent*<br>(N = 2) | *Early-only*<br>(N = 41 ) | *Persistent*<br>(N = 24 ) | P value |
| --- | --- | --- | --- | --- |
| Delivery Mode |  |  |  |  |
| Vaginal | 1 (50%) | 11 (27%) | 22 (92%) | < 0.001 |
| Pre-labor C-section | 1 (50%) | 15 (36.5%) | 2 (8%) |  |
| Post-labor C-section | 0 | 15 (36.5%) | 0 |  |
| Maternal race: White | 2 (100%) | 33 (80%) | 18 (75%) | 0.75 |
| BMI | 30 ± 1 | 27 ± 5 | 27 ± 7 | 0.82 |
| Chorioamnionitis | 0 | 6 (15%) | 4 (17%) | 0.81 |
| Meconium stained amniotic fluid | 0 | 9 (22%) | 7 (29%) | 0.58 |
| Birthweight | 4243 ± 605 | 3484 ± 479 | 3592 ± 372 | 0.06 |
| Infant antibiotics | 0 | 7 (17%) | 3 (13%) | 0.74 |
| Formula in hospital | 2 (100%) | 15 (37%) | 6 (25%) | 0.09 |
| Exclusive breastfeeding at week 2 | 1 (50%) | 22 (54%) | 15 (63%) | 0.77 |
| Duration of 2nd stage (median, IQR) | NA | 2.4 (1.15, 4.18) | 0.5 (0.32, 0.9) | 0.01 |
| Duration ROM (median, IQR) | 1.85 (0, 3.7) | 1.88 (0, 10.37) | 3.46 (0.94, 12.23) | 0.27 |

