## Supplemental Figures for "Delivery mode impacts newborn gut colonization efficiency"

### Supp Figure 1

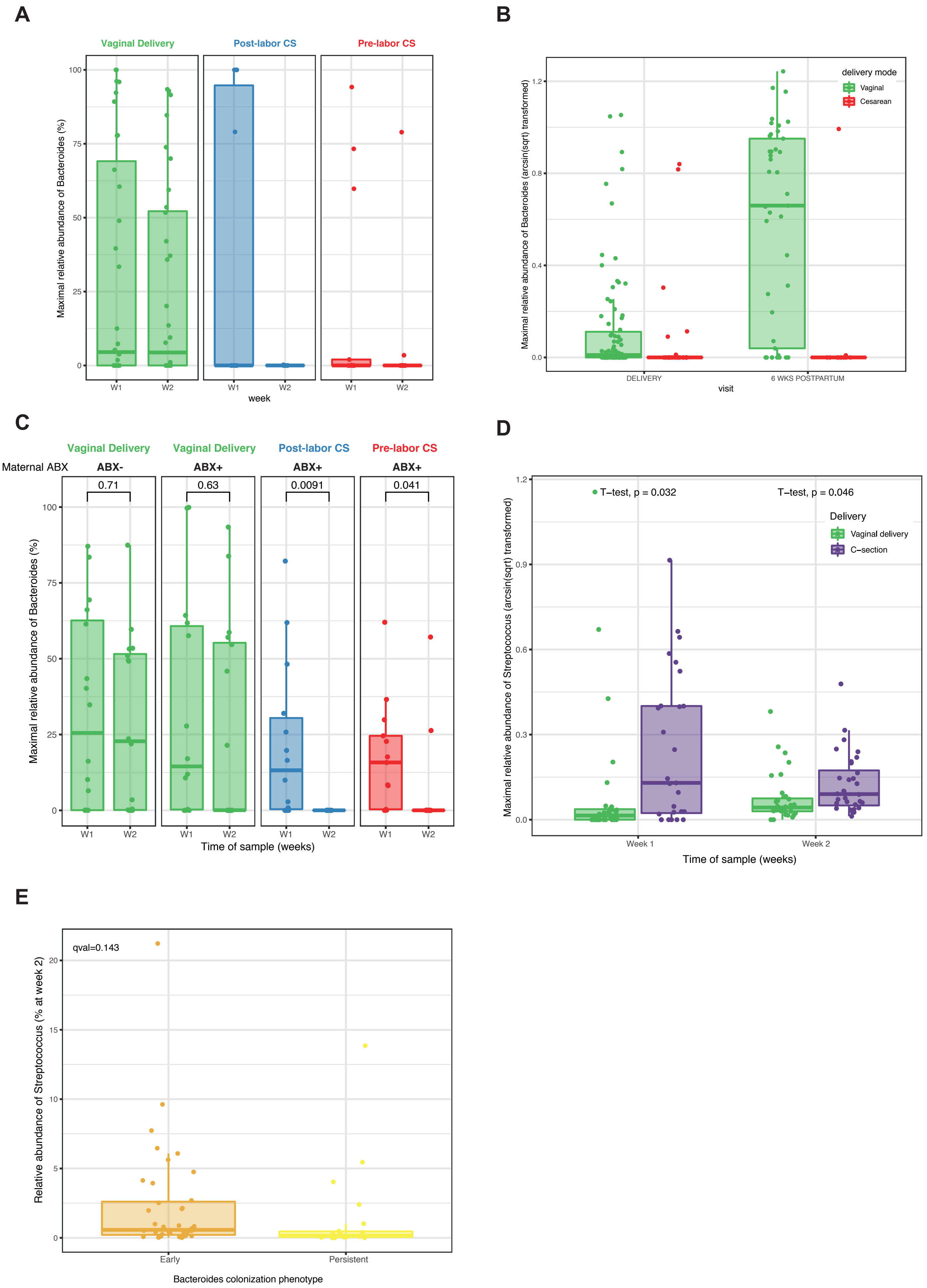

Supp Figure 2

**A**

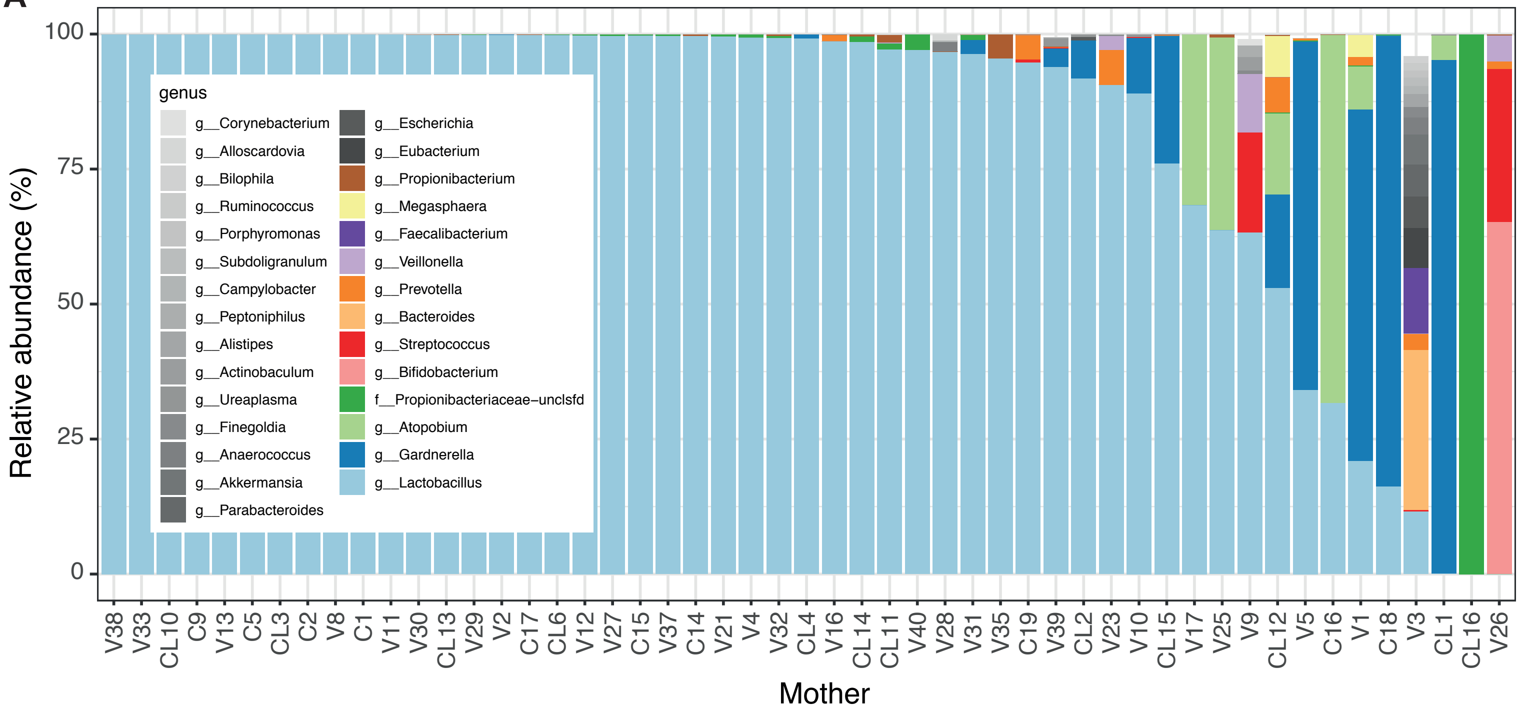


**B**

| | Maternal | Week 1 Child | Week 1 Shared | Week 2 Child | Week 2 Shared |
|---|---|---|---|---|---|
| **s__Lactobacillus_crispatus** | 29 | 2 | **1** (V29) | 0 | 0 |
| s__Lactobacillus_iners | 19 | 0 | 0 | 0 | 0 |
| s__Lactobacillus_jensenii | 13 | 0 | 0 | 0 | 0 |
| **s__Gardnerella_vaginalis** | 10 | 1 | **1** (CL15) | 1 | 0 |
| s__Lactobacillus_gasseri | 6 | 0 | 0 | 1 | 0 |
| **s__Atopobium_vaginae** | 6 | 1 | **1** (CL1) | 0 | 0 |
| s__Veillonella_atypica | 3 | 0 | 0 | 5 | 0 |
| s__Prevotella_bivia | 2 | 0 | 0 | 0 | 0 |
| s__Lactobacillus_sp_7_1_47FAA | 2 | 0 | 0 | 0 | 0 |
| **f__Propionibacteriaceae–unclsfd** | 2 | 11 | **1** (CL16) | 0 | 0 |
| s__Streptococcus_parasanguinis | 1 | 2 | 0 | 2 | 0 |
| s__Streptococcus_mitis_oralis_pneumoniae | 1 | 6 | 0 | 2 | 0 |
| s__Streptococcus_anginosus | 1 | 2 | 0 | 1 | 0 |
| s__Propionibacterium_acnes | 1 | 4 | 0 | 0 | 0 |
| s__Prevotella_timonensis | 1 | 0 | 0 | 0 | 0 |
| s__Parabacteroides_merdae | 1 | 4 | 0 | 0 | 0 |
| s__Faecalibacterium_prausnitzii | 1 | 4 | 0 | 0 | 0 |
| s__Eubacterium_rectale | 1 | 3 | 0 | 0 | 0 |
| s__Escherichia_coli | 1 | 18 | 0 | 14 | 0 |
| **s__Bifidobacterium_breve** | 1 | 3 | 0 | 11 | **1** (V26) |
| s__Bacteroides_vulgatus | 1 | 11 | 0 | 8 | 0 |
| s__Bacteroides_uniformis | 1 | 9 | 0 | 5 | 0 |
| s__Bacteroides_fragilis | 1 | 7 | 0 | 3 | 0 |

#families: 20, 10, 0
